# Effect of TIMPs and Their Minimally Engineered Variants in Blocking Invasion and Migration of Brain Cancer Cells

**DOI:** 10.1101/2024.06.05.597644

**Authors:** Elham Taheri, Maryam Raeeszadeh-Sarmazdeh

**Affiliations:** Chemical and Materials Engineering, University of Nevada, Reno

**Keywords:** TIMP minimal variants, glioblastoma multiforme (GBM), brain cancer, MMP inhibitors

## Abstract

Matrix metalloproteinases (MMPs) play a pivotal role in extracellular matrix (ECM) remodeling, influencing various aspects of cancer progression including migration, invasion, angiogenesis, and metastasis. Overexpression of MMPs, particularly MMP-2 and MMP-9, is notably pronounced in glioblastoma multiforme (GBM), a highly aggressive primary brain tumor characterized by diffuse and infiltrative behavior. Previous attempts to develop small molecule MMP inhibitors have failed in clinical trials, necessitating the exploration of more stable and selective alternatives. Tissue inhibitors of metalloproteinases (TIMPs), endogenous human proteins, offer promising potential due to their stability and broader interaction interfaces compared to small molecule inhibitors. In this study, we examined the effectiveness of wild-type human TIMP-1 and TIMP-3, alongside engineered minimal TIMP variants (mTC1 and mTC3), specifically designed for targeted MMP inhibition to reduce the migratory and invasive capabilities of GBM cells. Our investigation focused on these minimal TIMP variants, which provide enhanced tissue penetration and cellular uptake due to their small molecular weight, aiming to validate their potential as therapeutic agents. The results demonstrated that mTC1 and mTC3 effectively inhibit MMP activity, a critical factor in GBM aggressiveness, thereby highlighting their promise in controlling tumor spread. Given the lethality of GBM and the limited effectiveness of current treatments, the application of engineered TIMP variants represents a novel and potentially transformative therapeutic approach. By offering targeted MMP inhibition, these variants may significantly improve patient outcomes, providing new avenues for treatment and enhancing the survival and quality of life for patients with this devastating disease.

## 1. Introduction

Matrix metalloproteinases (MMPs) have a central role in extracellular matrix (ECM) remodeling (1). MMPs have several non-ECM substrates and contribute to several aspects of cancer progression, including migration, invasion, angiogenesis, and metastasis of tumors, by degrading the ECM or interacting with growth factors and cell receptors (2-6). MMP function is tightly regulated by other protease activators and inhibitors such as tissue inhibitors of metalloproteinases (TIMPs). Abnormal enzyme activity of specific MMPs was shown to have facilitated progression and invasion of glioblastoma multiforme (GBM), a highly aggressive primary brain tumor (7, 8). MMPs were also shown to contribute to disrupting the blood-brain barrier (BBB), which can result in neurodegenerative disorders (9) and potentially other brain-related diseases such as GBM (8, 10).

Specific MMPs, such as MMP-2/-9, are upregulated and associated with enhanced tumor invasion and therapeutic resistance in GBM (11-13). Further, overexpression of MMP-3, the endogenous activator of MMP-9, was associated with higher tumor grades (14, 15)(16, 17). MMPs ability to degrade the ECM facilitates the infiltration of tumor cells into surrounding brain tissue, contributing to the diffusive and infiltrative nature of GBM (18).

A more stable and selective MMP inhibitor alternative is needed because previous attempts at drug discovery to create small molecule MMP inhibitor therapeutics failed in clinical trials (19-22). A promising new direction for treatment development is the engineering of selective inhibitors based on tissue inhibitors of metalloproteinases (TIMPs), which are endogenous human proteins that compare to small chemical inhibitors and offer greater stability with a wider interaction interface (23, 24). Proteins such as TIMPs offer higher stability and selectivity and thus they have been considered as potential therapeutics for targeting MMPs. Further, TIMPs could be engineered to improve binding affinity and selectivity by targeting specific MMPs through protein engineering techniques, such as directed evolution and yeast surface display (23, 25). Recent efforts to engineer N-terminal domain of TIMP-2 using targeted evolution of a yeast displayed library resulted in potential protein scaffolds for selective targeting of MMPs (26-28). The four human paralogous genes encoding TIMPs 1 to 4 have a high level of sequence and structure homology, with a broad range of binding and inhibition to the MMP family of proteins. We have previously engineered minimal TIMP variants that inhibit MMPs effectively (29). These peptide-based TIMP variants provide more stability and higher selectivity compared to small molecules, and they also provide higher tissue and blood-brain barrier (BBB) penetration compared to proteins. Understanding the complex roles of MMPs in the ECM and the BBB is essential for developing targeted therapeutic strategies to inhibit MMP-mediated invasion and improve patient outcomes in this devastating disease. MMPs have also been shown to degrade the tight junctions of endothelial cells of BBB.

In this study, we examined the effect of wild-type human TIMP-1 and TIMP-3 recombinant proteins and previously engineered minimal TIMP variants (29), specifically designed to inhibit MMP-9, to reduce the migratory and invasive capabilities of GBM cells. MMP-9 is overexpressed in the GBM cell lines, T98G and A172 (30, 31). Our investigation was focused on two minimal TIMP variants, mTC1, and mTC3, which were developed to provide targeted inhibition of MMP activity—a crucial factor in the aggressive nature of GBM. By rigorously evaluating the efficacy of mTC1 and mTC3, we aimed to validate their potential as therapeutic agents and provide deeper insights into the MMP-TIMP dynamic within the GBM microenvironment. The engineered TIMP variants also provide higher tissue penetration and cellular uptake due to their small molecular weight (32, 33). Further, to improve the cellular uptake of these peptides, a cell-penetrating peptide (CPP) (34-36), due to of intracellular ability across the BBB and targeted delivery to GBM, was added at the N-terminus.

Given the lethality of GBM, the limited effectiveness of current treatment modalities, and the complexity of drug delivery across the BBB, advancements such as the application of minimal TIMP variants could revolutionize therapeutic approaches. Engineered minimal TIMP peptides offer new avenues for controlling tumor spread by targeting specific MMPs while having high drug deliverability to improve the survival and quality of life for patients battling this formidable brain tumor.

## 2. RESULTS

MMP-9 has been shown to be a key driver of GBM and its inhibition was shown to reduce the effects of GBM invasion in brain tumors (37, 38). We studied the effect of TIMP and its minimal variants on cell viability, cellular uptake, cell migration, and invasion in two GBM cell lines, T98 and A172. Both of these GBM cell lines have a high level of MMP-9 and MMP-3 expression (30, 31, 39-41). The engineered minimal TIMP variants (mTC1, mTC3), derived from DNA shuffling within the human TIMP family to generate a minimal TIMP hybrid library in yeast. Briefly, these peptides aimed to identify the dominant minimal MMP inhibitory regions. Subsequently, the synthesized minimal TIMP variants were screened against MMP-3 and MMP-9 using fluorescent-activated cell sorting (FACS) (29). Notably, several minimal TIMP variants, some comprising as few as 20 amino acids, were selected after screening against MMP-3cd or MMP-9cd and demonstrated either sustained or enhanced binding affinity to MMP-3 and MMP-9 (42).

### 2.1 TIMP and its variants do not affect GBM cell proliferation

The effect of TIMPs and their minimal variant peptides on cell proliferation were tested on GBM cell lines T98G and A172 alongside HeLa control cells using an MTT assay. Cells grown to confluency were treated with 0.02-2.5 μM of minimal TIMP variant mTC1. The viability of A172 cell lines persisted unaltered more than 90% until 1.25 μM of mTC1 then decreased to 88% at 2.5 μM (**Fig. 1**). The viability data for T98G cell line showed that mTC1 didn’t affect the cell proliferation, and viability remained above 90% until 0.32 μM and then decreased to 89% and 88% at 0.62 and 1.25 μM, respectively. The 2.5 μM of mTC1 reduced the viability of T98G cell line to 85%. The observed trend in the behavior of the HeLa cell line closely mirrored that of the GBM cell lines (**Fig. 1**). These findings demonstrate that mTC1 has little effect on viability in GBM and HeLa cell lines at lower concentrations, with only moderate reductions at higher concentrations, suggesting a potential for targeted therapeutic use with minimal cytotoxic effects at lower doses.

**Fig 1.**
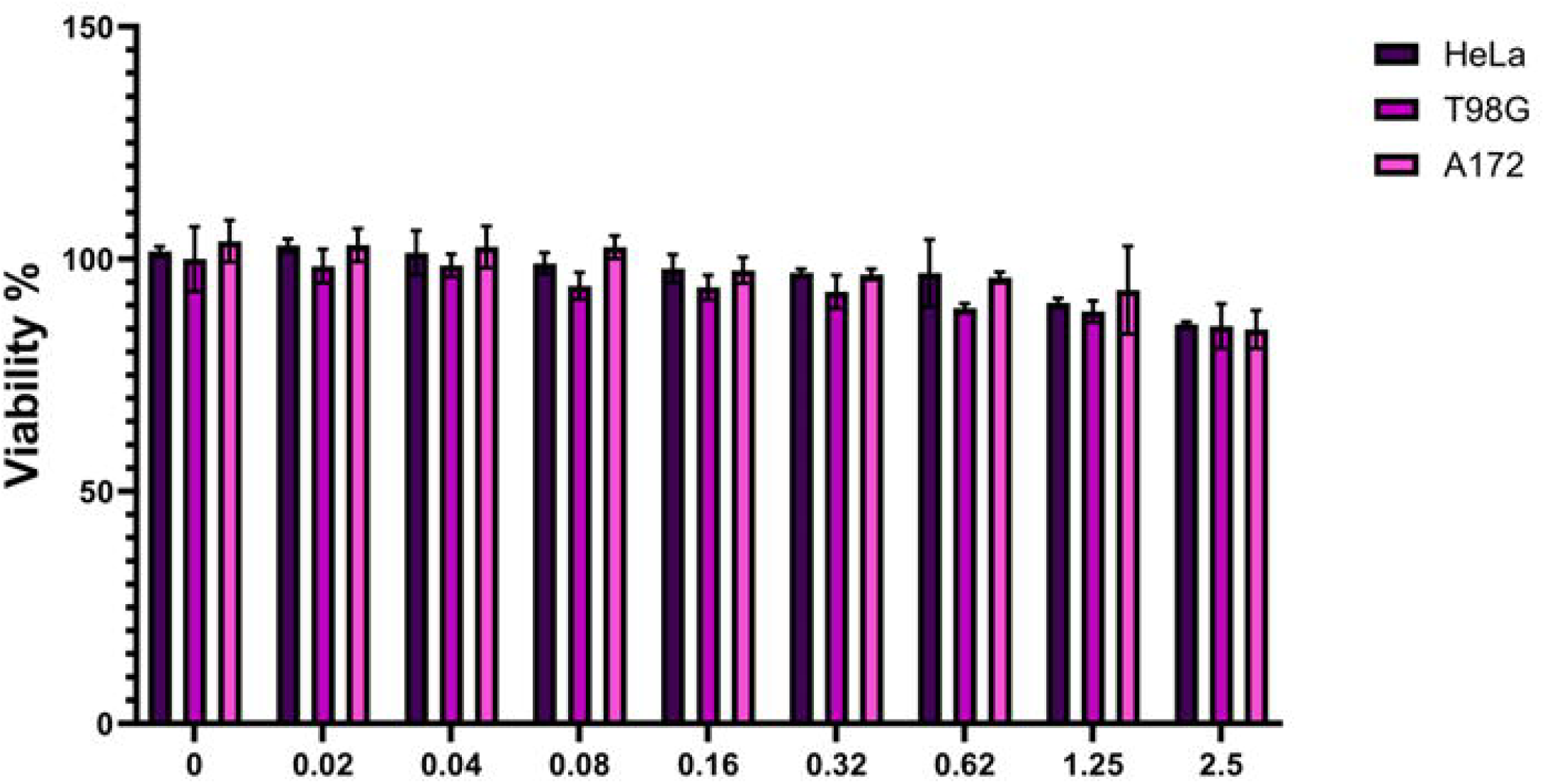
Effect of minimal TIMPs variants (mTC1) on the proliferation of GBM cell lines (T98G, A172) and HeLa cell line. The experiment was performed 24 h after adding mTC1 to GBM cell lines. MTT assay was performed as described in Methods for T98G and A172 and Hela cell lines as control. Bar graphs represent mean ± SEM of 4 different MTT experiments. Significance was obtained using One-way ANOVA. ^***^P < 0.001, ^**^P<0.01, ^*^P<0.05.

### 2.2. Cellular uptake

To improve cellular uptake of the minimal TIMPs variants, cell penetrating peptides (CPPs) were utilized, because of their ability to penetrate cells, become internalized, and break through various bio-barriers including the BBB (43-47). One of the most promising CPPs, TAT (RKKRRQRRR), has received a great deal of research due to its high cargo delivery efficiency (antibodies, nucleic acids, and nanoparticles) and effective biological barrier penetration (48, 49). We designed a new dual receptor recognizing cell penetrating peptide based on the combination of TAT and DGR to form (RKKRRQRRRDGR), a CendR (R/KXXR/K) motif, which can bind to NRP-1 (50), which is overexpressed receptor on tumor cells (50, 51), especially in GBM cell lines, including T98G and A172 (52) (53). Cellular uptake of a FAM-labeled CPP-conjugated peptide was evaluated on T98G and A172 cell lines using a ZOE fluorescence microscope. Fluorescence imaging demonstrated the uptake of FAM-labeled peptides in both GBM cell lines and localized in cytoplasm which was significantly higher than the uptake of mTC1 in both cell lines. Nuclei were visualized using Hoechst 10 mg/ml solution in water (1:2000 in PBS) (**Fig. 2**). This result confirmed peptide efficacy for intracellular delivery and this validation is crucial for establishing the peptide’s potential as a viable drug candidate.

**Fig 2.**
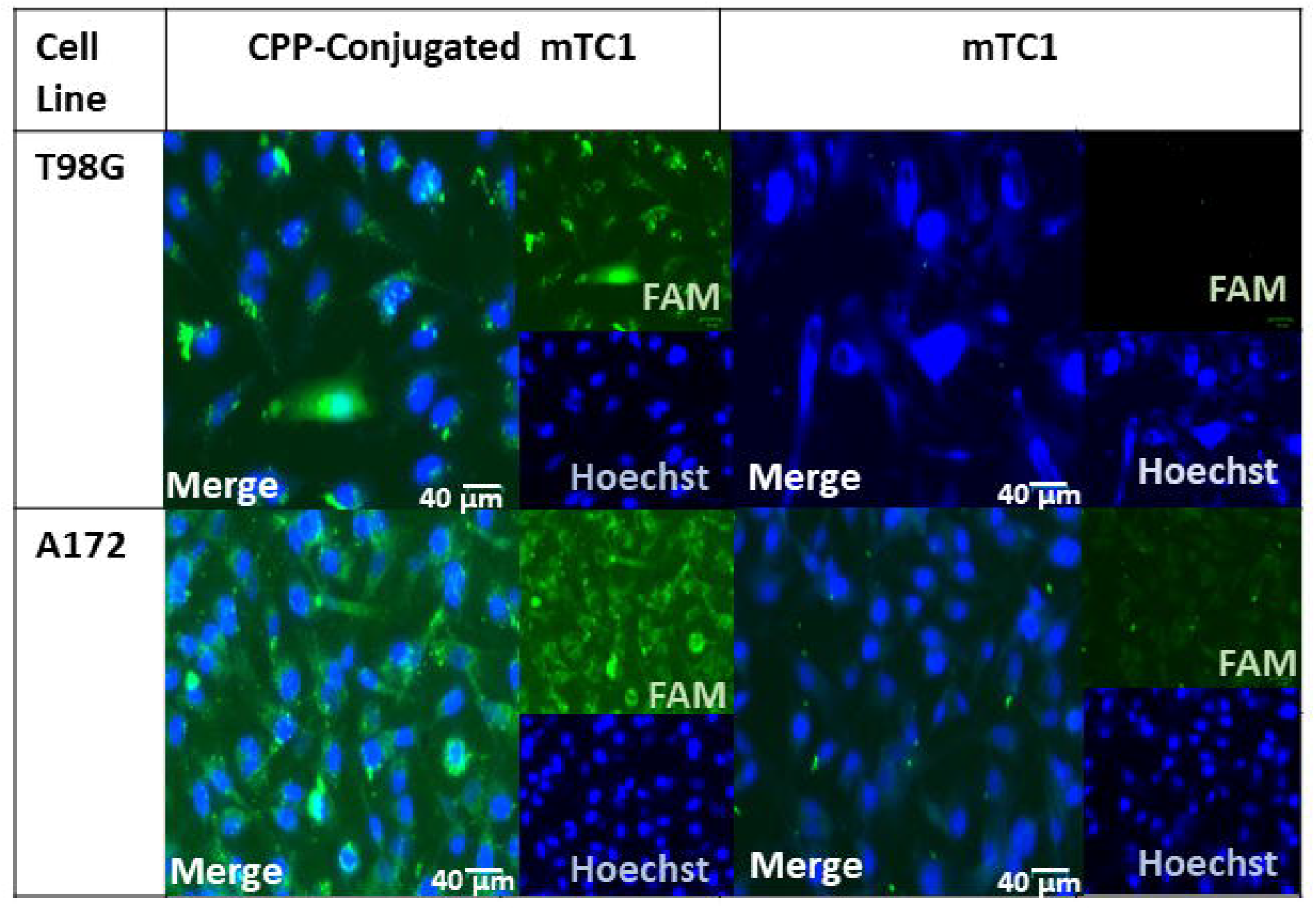
In vitro targeting of T98G and A172 cell lines. Fluorescence images of T98G and A172 cell lines after incubation with FAM-labeled peptides, mTC1, CPP-mTC1 for 4 h. T The images are the overlay of green and blue filters. Green, FAM-labeled peptide; blue, Hoechst-stained cell nucleus. The scale bars represent 40 μm.

### 2.3. TIMP and its minimal variants inhibit cell migration in GBM cells

A wound healing assay was employed to assess the impact of TIMPs and engineered TIMP variants on GBM cell migration using scratch assays. Our experimental findings demonstrate that treatment of T98G and A172 cells with varying concentrations (0.5 μM and 1.5 μM) of mTC1 and mTC3 effectively hindered their migratory behavior even better than TIMP-1 at the same concentration. Notably, after a 12 h incubation, mTC1 at concentrations of 0.5 μM and 1.5 μM led to a significant reduction in migrated cell numbers of the T98G cell line to 62% and 12%, normalized to without treatment, respectively (**Fig. 3B, S.1**.). Moreover, two concentrations of mTC1 showed similar effects on T98G migration after 18 h (**Fig. 3B, S.1**.). Additionally, we assessed the impact of mTC3 on migration inhibition of T98G. Our results indicate that after 12 h, mTC3 at a concentration of 1.5 μM, significantly reduced T98G migration to 25% and importantly, the inhibitory effect of mTC3 on migration remained consistent after 18 h, mirroring the observations at the 12 h mark (**Fig. 3B, S.1**.). To examine the impact of natural MMP inhibitors on T98G migration, we assessed two different concentrations (0.5 μM, 1.5 μM) of TIMP-1 and TIMP-3. TIMP-1 at 0.5 μM exhibited no significant effect after 12 h, while 1.5 μM of TIMP-1 reduced migration to 75%. Nevertheless, both 0.5 μM and 1.5 μM of TIMP-1 significantly reduced T98G cell migration after 18 h. Our findings indicate that following an 18 h incubation period, 0.5 μM and 1.5 μM TIMP-3 significantly decreased T98G migration to 65% and 25%, respectively; however, after 12 h, 0.5 μM did not exhibit a significant effect on migration (**Fig. 3B, S.1**.). We also evaluated the effect of TIMPs and engineered TIMP variants on A172 migration. 1.5 μM mTC1 significantly reduced A172 migrated cells, to 20% and 40%, after 12 and 18 h, respectively, while 0.5 μM mTC1 didn’t show a significant effect after 18 h (**Fig. 4B, S.2**.). Moreover, 1.5 μM TIMP-1 reduced A172 migrated cells to 75% and 65% after 12 and 18 h, respectively (**Fig. 4B, S.2**.). After 12 h, both 0.5 and 1.5 μM mTC3 significantly reduced migrated cell to 45% and 35%, respectively (**Fig. 4B, S.2**.). These results suggest that the engineered minimal TIMP variants, particularly mTC1 and mTC3, exhibit a strong potential for inhibiting GBM cell migration more effectively than natural TIMP-1, and this substantial reduction in the migration of T98G and A172 cells highlights their efficacy as potential therapeutic agents for GBM.

**Figure 3.**
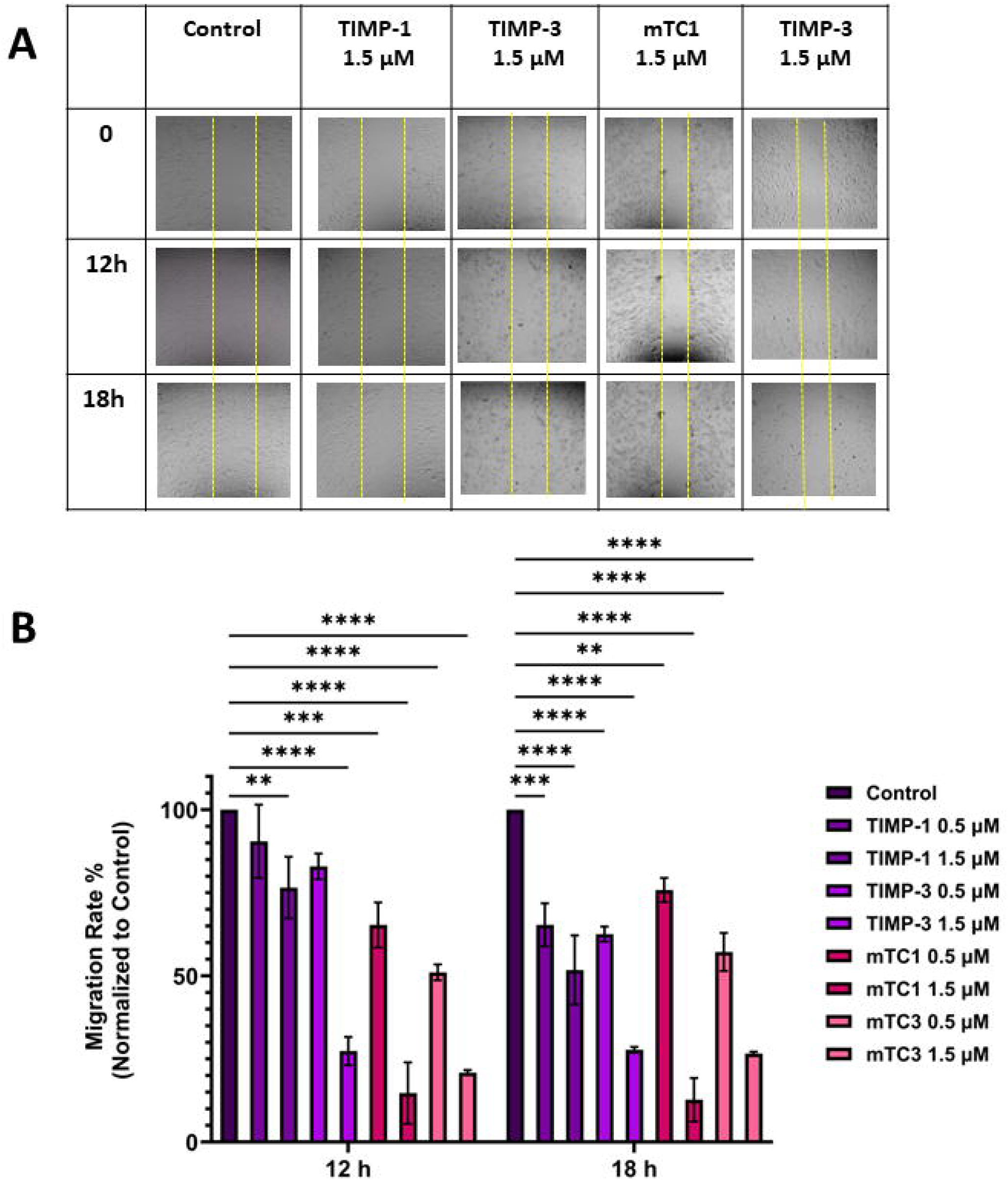
Effect of TIMPs and minimal TIMPs variants on the migration of T98G cell line. (A) Representative of migration T98G in the presence and absence of TIMPs and minimal TIMPs variants. The cells were visualized by light microscopy using a 10x magnification lens on 0, 12 h, and 18 h, after adding 1.5 μM of TIMPs (TIMP-1, TIMP-3) and minimal TIMPs variants (mTC1, mTC3), (B) Calculated fold of migration of T98G cell line. The created wounds were analyzed using Image J software and wound closure percentage was calculated (The data were normalized to control). The experiment was repeated two times; means and standard error are shown. ^***^P < 0.001, ^**^P<0.01, ^*^P<0.05 as determined by a t-test comparing inhibition in the presence of the various inhibitors versus the untreated control.

**Figure 4.**
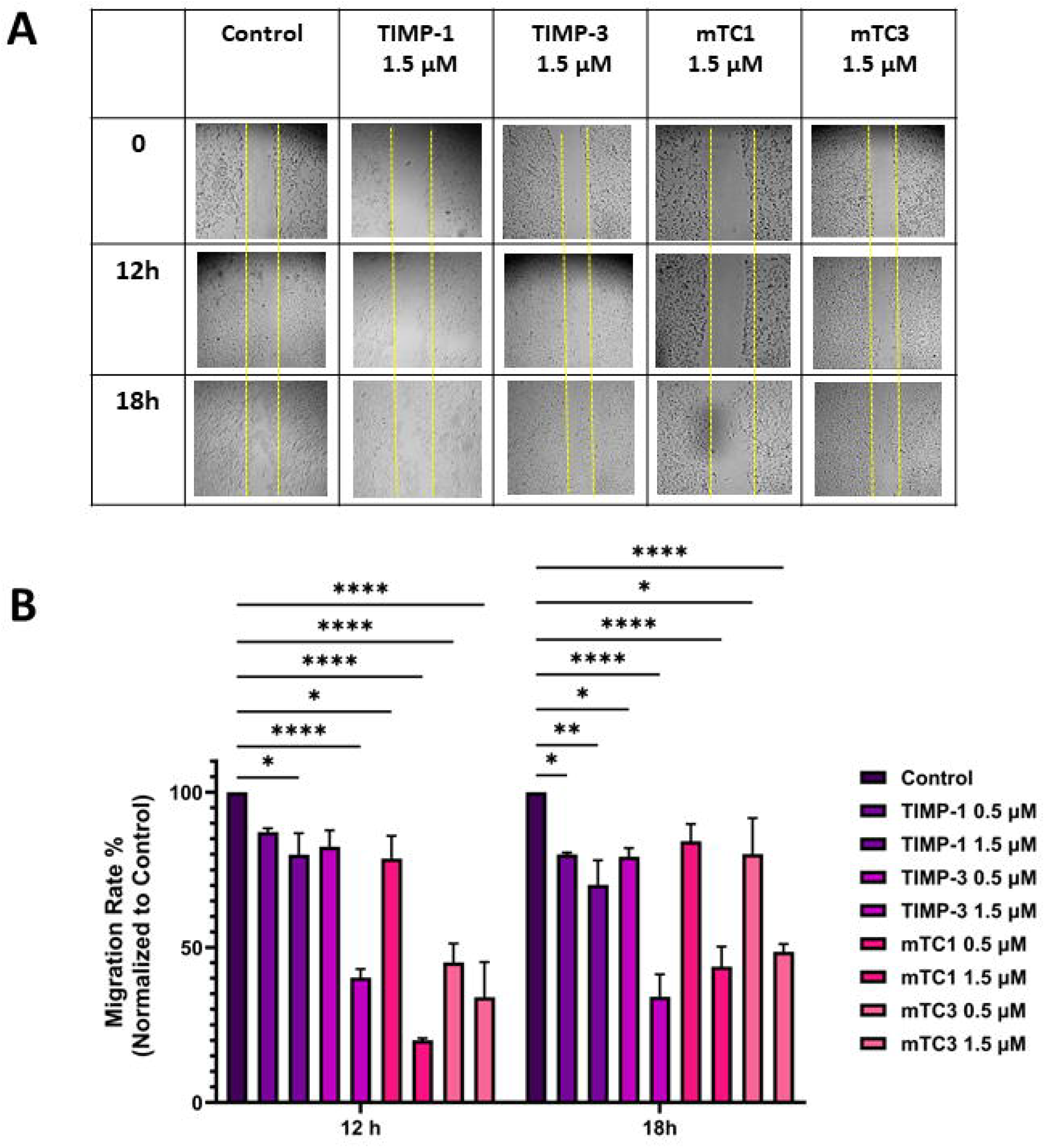
Effect of TIMPs and minimal TIMPs variants on the migration of A172 cell line. (A) Representative of migration T98G in the presence and absence of TIMPs and minimal TIMPs variants. The cells were visualized by light microscopy using a 10x magnification lens on 0, 12 h, and 18 h, after adding 0.5 μM and 1.5 μM of TIMPs (TIMP-1, TIMP-3) and minimal TIMPs variants (mTC1, mTC3), (B) Calculated fold of migration of T98G cell line. The created wounds were analyzed using Image J software and wound closure percentage was calculated (The data were normalized to control). The experiment was repeated two times; means and standard error are shown. ^***^P < 0.001, **P<0.01, ^*^P<0.05 as determined by a t-test comparing inhibition in the presence of the various inhibitors versus the untreated control.

### 2.4. Invasion tests using TIMP and engineered variants

To study the impact of MMP inhibition on the invasive properties of T98G and A172 cells, Matrigel Transwell assays were performed. The GBM cell lines T98G and A172 were grown on 24-well Matrigel Transwells and treated with vehicle, 0.5 μM or 1.5 of μM TIMP protein or its minimal variants. While TIMP-1 had a minimal inhibitory effect on blocking the invasion of GBM cells, the number of invasive GBM cells that traversed the Matrigel significantly decreased following mTC1 treatment compared to the control cells in a dose-dependent manner with 1.5 μM being the most effective. After 20 h, T98G cell invasion decreased by 30% and 18% with mTC1 concentrations of 0.5 μM and 1.5 μM, respectively compared to TIMP-1 with 39% and 38%, for concentrations of 0.5 μM and 1.5 μM, respectively (**Fig. 5B**). To further, we investigated the effect of mTC1 on the invasion of the A172 cell line. This further proves the great therapeutic potential of the minimal TIMPs (**Fig. 6B**). Alongside, we assessed the potential of minimal TIMP and TIMP-1 to inhibit invasion in the A172 cell line. The observed trend of invasion inhibition in the A172 cell line was consistent with that of the T98G cell line.

**Figure 5.**
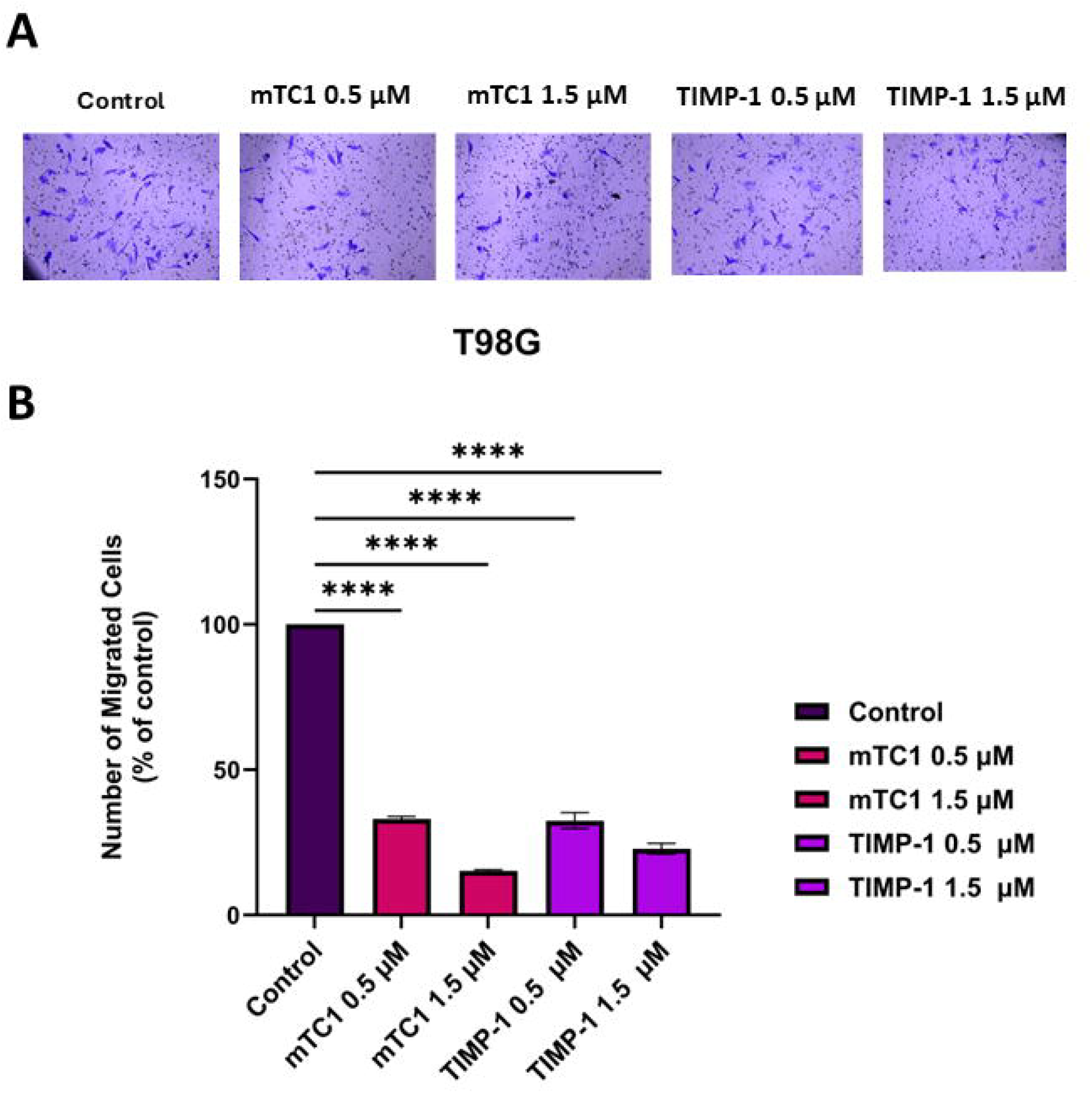
Effect of TIMP-1 and minimal TIMPs variant (mTC1) on the T98G cell line. (A) Representative of invasion T98G cell line in the presence and absence of TIMP-1 and mTC-1. The cells were fixed in 100% methanol stained with 2% crystal violet and then visualized by light microscopy using an X10 magnification lens. (B) Calculate migration of T98G. The cells were counted using ImageJ software and normalized to the count of untreated cells. Significance was obtained using One-way ANOVA. ^*^p < 0.05, ^***^p < 0.001, ^****^p < 0.0001.

**Figure 6.**
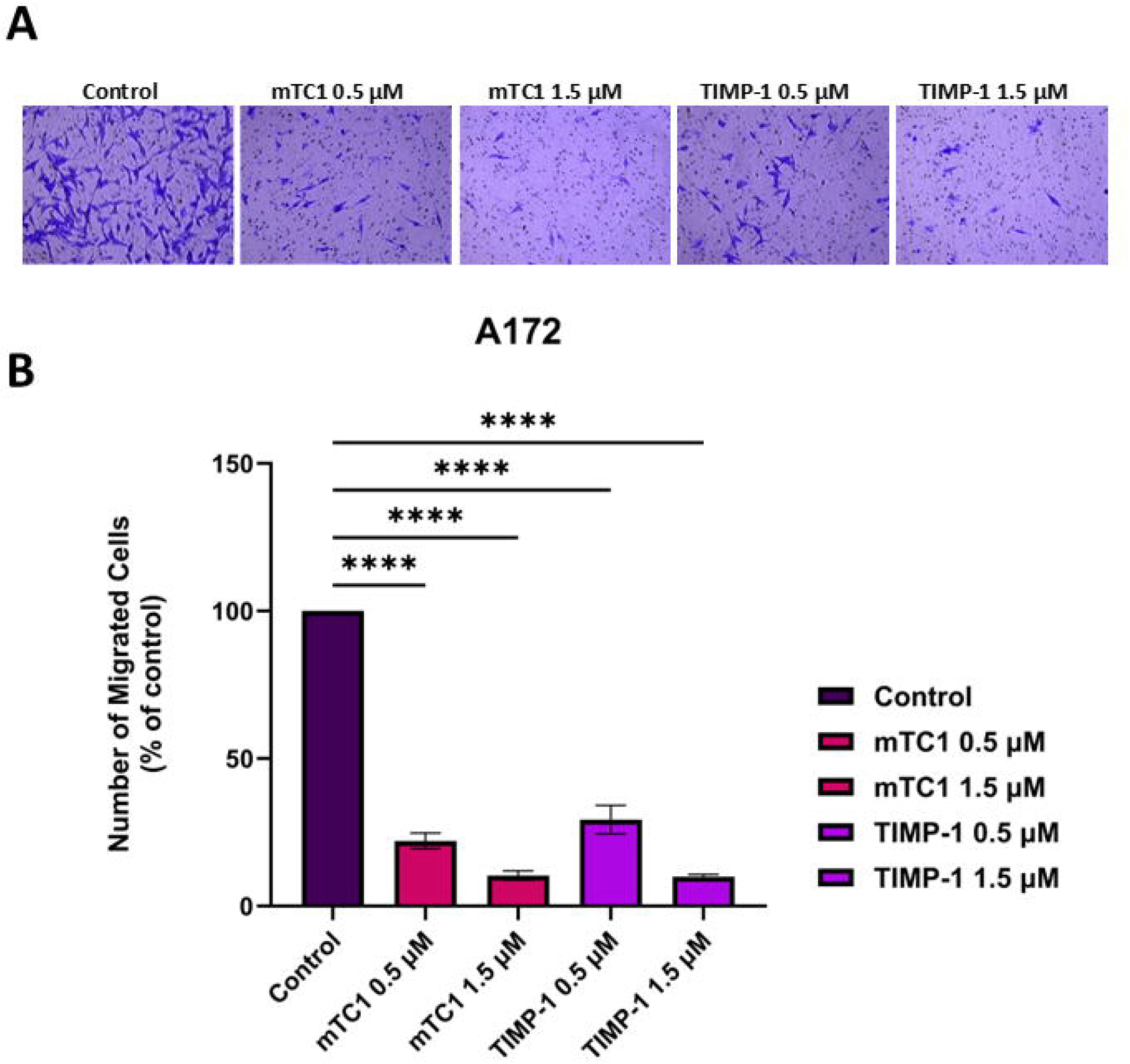
Effect of TIMP-1 and minimal TIMPs variant (mTC1) on the A172 cell line. (A) Representative of invasion T98G cell line in the presence and absence of TIMP-1 and mTC-1. The cells were fixed in 100% methanol and stained with 2% crystal violet and then visualized by light microscopy using a X10 magnification lens. (B) Calculate fold of migration of T98G. The cells were counted using ImageJ software and normalized to the count of untreated cells. Significance was obtained using One-way ANOVA. ^*^p < 0.05, ^***^p < 0.001, ^****^p < 0.0001.

## 3. Material and methods

### 3.1. Chemical Reagents

Eagle’s minimum essential medium (EMEM) Transwell with 0.8 μm pore polycarbonate membrane inserts, and Matrigel basement membrane matrix, LDEV-free were obtained from Corning (Corning, Glendale, Az, USA). Dulbecco’s modified Eagle’s medium (DMEM), 0.05% trypsin/0.02% EDTA, DMSO (Dimethyl sulfoxide), MTT (3-(4,5-dimethylthiazol-2-yl)-2,5-diphenyltetrazolium bromide), Hoechst 33342, trihydrochloride trihydrate, 10 mg/mL were purchased from Life Technologies (Thermo Fisher, Waltham, MA, USA). Dulbecco’s phosphate-buffered saline (DPBS), fetal bovine serum (FBS), and 100X antibiotics (10,000 U/mL of penicillin and 10,000 μg/mL streptomycin) were obtained from Gibco (Gibco, USA). All peptides including minimal TIMPs variants, CPP-conjugated minimal TIMP (mTC1) and 5□carboxyfluorescein (FAM) labeled peptides (mTC1, CPP, CPP-conjugated mTC1) were synthesized by the Genscript Inc. (Piscataway, NJ, USA).

### 3.2. Cell Culture

The T98G glioma cell line (CRL-1690™, ATCC, Manassas, VA, USA) and the HeLa cell line (CCL-2 ™) were generously provided as a gift by Dr. Vincent Lombardi (Pharmacology, University of Nevada, Reno) and Dr. Bahram Parvin (Biomedical Engineering, University of Nevada, Reno), respectively. A172 glioma cell line was purchased from ATCC (CRL-1620 ™, ATCC, Manassas, VA, USA). T98G and HeLa cell lines were cultured in EMEM media supplemented with 10% FBS and 1% of the antibiotic mixture (penicillin, 100 IU/mL and streptomycin, 100 μg/mL) in T-75 flasks (Thermo Fisher, USA). A172 cell line was cultured in high glucose DMEM media plus 4 mM L-glutamate and 1 mM sodium pyruvate (Thermo Fisher) supplemented with 10% FBS and 1% penicillin and streptomycin mixture. The cells were incubated at 37 °C in the presence of 5% CO_2_. The medium was changed every third day, and the cells were passaged at approximately 80% confluence. For cell sub-culturing, the cells were washed with 5 mL DPBS, then 2 mL of 0.05% trypsin/0.02% EDTA were added to flask and incubated for 3 min until the cells detached and fresh culture medium was added, and dispensed into new culture flasks.

### 3.3. Minimal TIMPs amino acid sequences

The minimal TIMPs peptides, mTC1 (CTCVPPHPQTAFLCTWQSLRSQIA) and mTC3 (CTCVPPHPQTAFLCTWQSLRSQIA ), hybrid of TIMP-1/TIMP-3 and TIMP-2/TIMP-3 sequences (42), respectively were identified based on their high affinity to MMPs and then their inhibitory effect for GBM was investigated. All minimal TIMPs peptides candidates employed in this investigation were synthesized by Genscript peptide services (Genscript USA, New Jersey). Genscript confirmed that all peptides were obtained with >90% purity. Before use, all peptides were dissolved in the 5 μl of formic acid to form 300 mM concentration, then they were diluted in TNC buffer (50 mM Tris HCl, pH 7.5, 150 mM NaCl, 10 mM CaCl_2_) to the final concentration of 1.5 μM for testing in the GBM cell lines. Peptides that were reconstituted were kept at -20 °C until needed.

### 3.4. Cellular uptake

The cellular uptake of fluorescence-labeled peptide was evaluated on T98G and A172 cell lines. For quantitative analysis, 5 × 10^5^ cells per well of each cell line were plated in fibronectin-coated (2 μg/cm^2^) 35 mm glass base dishes (Thermo Scientific, Rochester, NY, USA). After incubation for 24 h and reaching 90% confluency, 1.5 μM of fluorescence-labeled peptide were added into the glass base dishes and incubated for 4 h 37°C and 5% CO_2_. Then the cells were washed twice with PBS, trypsinized, resuspended in 0.5 mL PBS and finally analyzed by a flow cytometer (BD Accuri C6 Flow Cytometer, BD Biosciences).

For the qualitative analysis, 1 × 10^5^ cells of T98G and A172 cell lines were seeded on 35 mm glass base dishes (Thermo Scientific, Rochester, NY, USA) coated with 2 μg/cm^2^ fibronectin and allowed to grow for 24 h. After another 4 h incubation with 1.5 μM of fluorescence-labeled peptide, cells were washed twice with cold PBS, fixed with 4% paraformaldehyde for 30 min in room temperature, then stained with Hoechst (10 mg/ml solution in water, diluting the Hoechst stock solution 1:2,000 in PBS) for 5 min. Finally, the cells were imaged using a fluorescence microscopy (ZOE™ Fluorescence Imaging, Bio-Rad, Hercules, CA, USA).

### 3.5. Wound Healing Assay

The wound healing assay was used to investigate the effects of minimal TIMP variants on GBM cell migration dynamics. T98G and A172 cells were grown to approximately 80% confluency, then, 1 × 10^6^ cells of each were plated in 12-well plates containing 1 mL of appropriate medium. After 24 h incubation at 37 C and 5% CO_2_, and reaching 80% confluency, straight scratch wounds were made in the middle of confluent cells in each well using a sterile P200 pipette tip along the diameter of the well. The cells were washed with Dulbecco’s phosphate-buffered saline (DPBS) (pH 7.4) to remove the debris and floating cells. After removing DPBS, cells were incubated in a medium with or without different concentrations (0.5 μM and 1.5 μM) of TIMP-1 or minimal TIMP variants for 12 and 18 h. The wounded areas were examined, and an image was taken under a light microscope (Inverted Microscope, Fisher Scientific, USA) at 10X magnification in the zone to view cell migration at different time points (0, 12, and 18 h). The experiments were performed in duplicate. The closure of the wounded area was calculated by the ImageJ software (National Institutes of Health, Bethesda, MD, USA). The migration distance or the percentage of wound closure was calculated using the following equation (54):

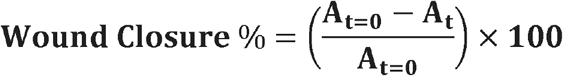

Where, A_t=0_ is the initial wound area, A_t_ is the wound area after hours of the initial scratch, both in μm^2^.

### 3.6. Matrigel Invasion Assay

In vitro Matrigel invasion assay was performed in a Transwell culture chamber system. The filter membranes with 8 μm pores and 0.33 cm^2^ were coated with 100 μL of 0.250 mg/mL Matrigel (BD Biosciences, USA) at 37°C / 5% CO_2_ for 2 h. Then, 2.5×10^4^ GBM cells (T98G, A172) were resuspended in 500 μL of serum-free EMEM medium in the presence or absence of minimal TIMPs and added to the upper chamber. The lower chamber was filled with 0.75 mL of complete medium (10% FBS) as chemo-attractant and cells were then incubated at 37°C / 5% CO_2_ for 24 h. After incubation, non-migrated cells on the upper surface of the membrane were removed using sterilized cotton swabs, migrated cells on the lower surface were fixed in 100% methanol and stained with 2% crystal violet. Multiple fields of cells were counted randomly in each well under a light microscope (Fisher brand™ Entry Level Research Grade Inverted Microscope, Fisher Scientific, USA) at 10X magnification. Data were expressed as the percentage of invasive cells as compared with the control (55).

### 3.7. MTT Assay

The MTT (3-(4,5-dimethylthiazol-2-yl)-2,5-diphenyltetrazolium bromide) assay was performed to assess the viability of T98G/A172 and HeLa cells treated with different concentration of TIMP and minimal TIMP variants. Cells were seeded in 96-well plates and allowed to adhere overnight at 37°C and 5% CO_2_. The cells were then treated with various concentrations of TIMP, minimal TIMP proteins, media, and buffer as control solutions for 24 h while incubated at 37°C and 5% CO_2_. After treatment, 20 μL MTT solution (0.5 mg/mL) was added to each well and incubated for 4 h at 37°C with 5% CO_2_. Viable cells contain NAD(P)H-dependent oxidoreductase enzymes and reduce the MTT to formazan. Formazan crystals were dissolved using a 100 μL of dimethyl sulfoxide (DMSO) as a solubilization solution. The color intensity of the resulted solution is quantified by measuring absorbance at 570 nm wavelength.

### 3.8. Statistical Analysis

All experiments were performed in triplicate, and data were expressed as mean ± standard deviation (SD). Statistical analysis was performed using GraphPad Prism software. One-way analysis of variance (ANOVA) followed by Tukey’s post hoc test was used to determine the significance of differences between multiple groups. A p-value <0.05 was considered statistically significant.

## 4. DISCUSSION

GBM is an aggressive brain cancer with unique challenges in developing therapeutics from difficulties in reaching medications across the blood-brain barrier (BBB). Specific MMPs such as MMP-9 have been shown to play a key role in both migration and invasion of GBM tumor cells, and disturbing the BBB integrity, contributing to the progression of the disease. To overcome the challenges in finding efficient treatments for GBM, we investigated the treatment of GBM cell lines, T98G and A172, where MMP-9 is overexpressed, using engineered minimal TIMP variants generated using DNA shuffling, directed evolution, and yeast surface display.Since these engineered minimal TIMPs showed better stability compared to small molecules and higher tissue and blood-brain barrier (BBB) penetration compared to the protein, we considered as ideal therapeutic candidates for MMP inhibition in GBM cell lines with higher levels of MMPs’ expression.

Our results indicated that migration and invasion of GBM cell lines, T98G and A172, can be modulated by blocking MMP activity, and using an effective MMP-3/9 inhibitor can reduce migration and invasion in these cell lines. Using wound healing assay, our results demonstrated that the studied engineered minimal TIMPs, mTC1 and mTC3 peptides, effectively reduced migration at both concentrations. Interestingly, 1.5 μM of mTC1 and mTC3 reduced the migration of T98G cells to 10% and 25%, respectively. Moreover, these engineered minimal TIMPs exhibit a more pronounced inhibitory effect on migration compared to natural TIMPs (TMP-1 and TIMP-3). Transwell Matrigel assay for testing the effect of mTC1 on the invasion of GBM cell lines, as a better migratory inhibitor in both cell lines, confirmed the ability of mTC1 to inhibit the invasion of both GBM cell lines. The minimal cytotoxicity observed at lower concentrations indicates a promising therapeutic index, making these variants viable candidates for further development and clinical application. Additionally, by integrating cell-penetrating peptides (CPPs) to the minimal TIMP peptides, we achieved enhanced intracellular delivery, ensuring that these engineered variants effectively penetrate the BBB and localize within the tumor cells.

In summary, this study provides robust evidence supporting the use of minimal TIMP variants as a novel therapeutic approach for GBM. The ability of mTC1 and mTC3 to inhibit MMP-mediated invasion and migration paves the way for future research to optimize these variants for clinical use. Future directions should include in vivo studies to assess the long-term efficacy and safety of these peptides, and the exploration of the development of advanced delivery systems to improve targeting and penetration. By addressing the invasive and resistant nature of GBM, these innovative treatments hold the potential to significantly improve patient survival and quality of life, marking a pivotal step forward in the fight against this aggressive brain tumor.

## ABBREVIATIONS

MMP: matrix metalloproteinase
FACS: fluorescent-activated cell sorting
scFv: single-chain antibody fragment
TIMP: tissue inhibitors of metalloproteinases
GBM: glioblastoma multiforme
CPP: cell-penetrating peptide.

## Author contributions

M.R.-S., E.T; conception, E.T.; performed the experiments, E.T., M.R.-S., data analysis, writing, and editing the manuscript drafts. M.R.-S. funding and supervision.

## Acknowledgment

We would like to thank Dr. Bahram Parvin (Biomedical Engineering-UNR), and Dr. Vincent Lombardi (Pharmacology-UNR) for the generous gift of the GBM and HeLa cell lines.

