## Supplementary figures and images for "Effect of TIMPs and Their Minimally Engineered Variants in Blocking Invasion and Migration of Brain Cancer Cells"

### Supplemental Data 1

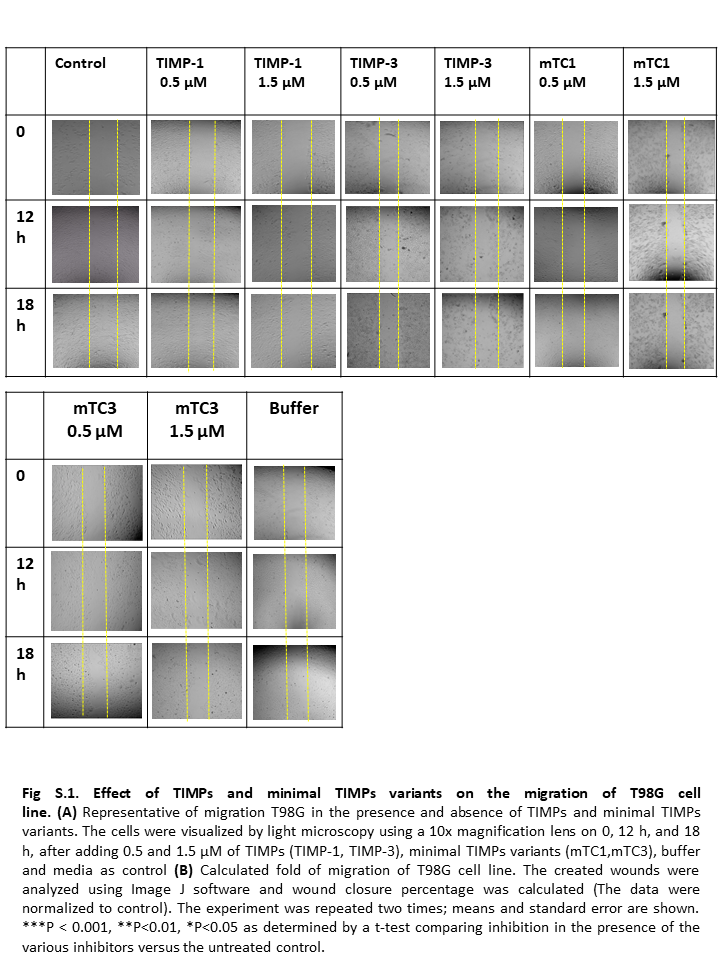

### Supplemental Data 2

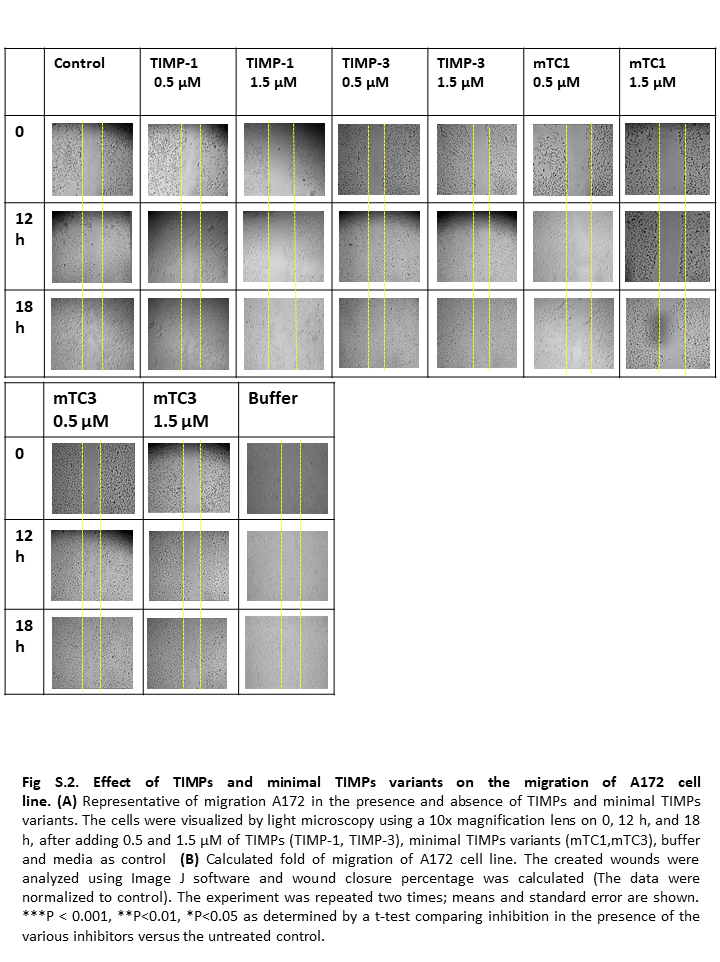
